# The proteome of small urinary extracellular vesicles after kidney transplantation as an indicator of renal cellular biology and a source for markers predicting outcome

**DOI:** 10.1101/845941

**Authors:** Fabian Braun, Markus Rinschen, Denise Buchner, Katrin Bohl, Martin Späth, Heike Göbel, Corinna Klein, Oliver Kretz, Victor G. Puelles, Daniel Bachurski, Ingo Plagmann, Roger Wahba, Michael Hallek, Bernhard Schermer, Thomas Benzing, Tobias B. Huber, Andreas Beyer, Dirk Stippel, Christine E. Kurschat, Roman-Ulrich Müller

## Abstract

Kidney transplantation is the preferred renal replacement therapy available. Yet, the biological processes during and after kidney transplantation and how they translate into the overall functional graft outcome are insufficiently understood. Recent developments in the field of extracellular vesicle research allow the deeper exploitation of this non-invasive source. We separated small urinary extracellular vesicles (suEVs) throughout the course of living donor kidney transplantation. SuEVs were collected longitudinally from both the donor and the recipient in 22 living donor kidney transplantations. Unbiased proteomic analysis revealed specific temporal patterns of suEV proteins indicative of the cellular processes involved in the allograft’s response after transplantation with proteins playing a role in complement activation being among the most dynamically regulated components. Using a leave-one-out cross validation model, we identified potential prognostic markers of kidney function at 1 year after transplantation. One of the proteins identified – phosphoenol pyruvate carboxykinase (PCK2) – could be confirmed in an independent validation cohort of another 22 donor-recipient pairs using targeted mass spectrometry. This study sheds the light on early molecular processes during the course of kidney transplantation and shows the future potential of suEVs as a source of biomarkers in this setting. The data set is provided as a unique resource directly accessible through an online tool that allows dynamic interrogation of this first comprising suEV proteome atlas after kidney transplantation.

**One Sentence Summary:** This study represents the first atlas of the proteomic changes in small urinary extracellular vesicles throughout living donor kidney transplantation identifying PCK2 abundance as a biomarker for renal function 12 months after transplantation

## Introduction

Chronic kidney disease has emerged as one of the major health problems and cost factors for health care systems worldwide affecting all age groups (*1*). While renal replacement therapy by dialysis can reinstate kidney function regarding acid-base homeostasis, water and toxin elimination to a certain extent, it is accompanied by a dramatic increase in both cardiovascular and overall mortality (*2, 3*). The treatment strategy that results in increased quality of life and patient survival is kidney transplantation (*4, 5*). However, the number of potential recipients outweighs the number of donors considerably. This unmet need has prompted a rise in living donor transplantation circumventing the need for dialysis (*6*).

Yet, transplant rejection and chronic allograft nephropathy limit organ survival making repetitive transplantation necessary in many cases (*7*). Diagnosis of the underlying disorder relies primarily on renal biopsy via hollow needle tissue sampling involving both a bleeding risk with possible organ damage and the potential of misdiagnosis due to sampling errors (*8, 9*). With primary urine passing all cells of the nephrons, the identification of urinary markers would be of great potential to complement histology. To date, urinary sediment and proteinuria are still the only tools available in clinical routine (*10*).

Recent years have shifted different extracellular vesicles (EVs) into the research focus for both their biological function as well as their potential as a source of biomarkers for various diseases (*11*). In contrast to analyses from full urine, EVs – being actively secreted by nephron-lining epithelial cells – hold the promise to reflect cellular biology more directly. Most studies have discriminated between three types of EVs (exosomes, microvesicles and apoptotic bodies) depending on their biogenesis (*12, 13*).

Current investigations, however, have depicted both the shortcomings of past separation methods and vesicle characterizations as well as a considerable overlap in both size, density as well as protein and nucleic acid content of all three groups (*14–18*). As a consequence, the International Society of Extracellular vesicles (ISEV) proposes an operational terminology for separated extracellular vesicles, indicating the biochemical and physical properties (*15*).

To quantitatively assess the proteome of small urinary EVs (suEVs) and its changes throughout living donor transplantation, we employed a simple separation protocol for urinary EVs through differential (ultra-)centrifugation. The separated suEVs were analyzed for their protein content by mass spectrometry using label-free quantification. The resulting data set represents a novel and unique resource giving insight into biological processes involved throughout renal transplantation. Additionally, correlation with kidney function after transplantation in combination with targeted mass spectrometry for specific markers in a validation cohort provides a first insight regarding the power of this approach for outcome prediction after kidney transplantation.

## Results

### A fast and widely applicable protocol of differential centrifugation yields a robust separation of suEVs

We established a fast, cost-efficient and easy protocol for the separation of small urinary extracellular vesicles using differential ultracentrifugation to allow for a potential transfer to clinical routine in the future (Fig. 1A). After the addition of preservatives and protease inhibitors, a centrifugation step of 17.000 g performed at 4°C for 20 minutes pellets all cells, cellular debris and large to medium EVs. Analysis of the final suEV pellet via transmission electron microscopy (TEM) and Nanoparticle Tracking analysis via ZetaView depicted a wide variety of small EVs with a typical cup-shaped morphology ranging from suEVs of 30 – 60 nm to a small fraction of large urinary EVs with a diameter up to 500 nm in diameter (Fig. 1B/C and S1A). To validate the stability of suEV proteins in long-term storage after dissolving the final pellet in 8 M urea buffer, we performed western blot analyses after 1 and 6 months at −80°C for small EV marker proteins and detected no storage-dependent differences in immunoblot signal (Fig. 1D). Additionally, our separation protocol achieved a decrease in the ratio of the ubiquitous urinary protein uromodulin to the suEV protein TSG101 (Fig. S1B) and we were able to robustly detect the tetraspanins CD9, CD63 and CD81 – typically present on the surface of EVs – in a flow cytometric bead assay (Fig. S1C) of the final suEV pellet. A pilot mass spectrometry analysis confirmed that our protocol separated particles containing marker proteins for all segments of the nephron (*19*) as well as Alix and TSG101 – two proteins typically found in EVs (Fig. 1E).

**Fig.1:**
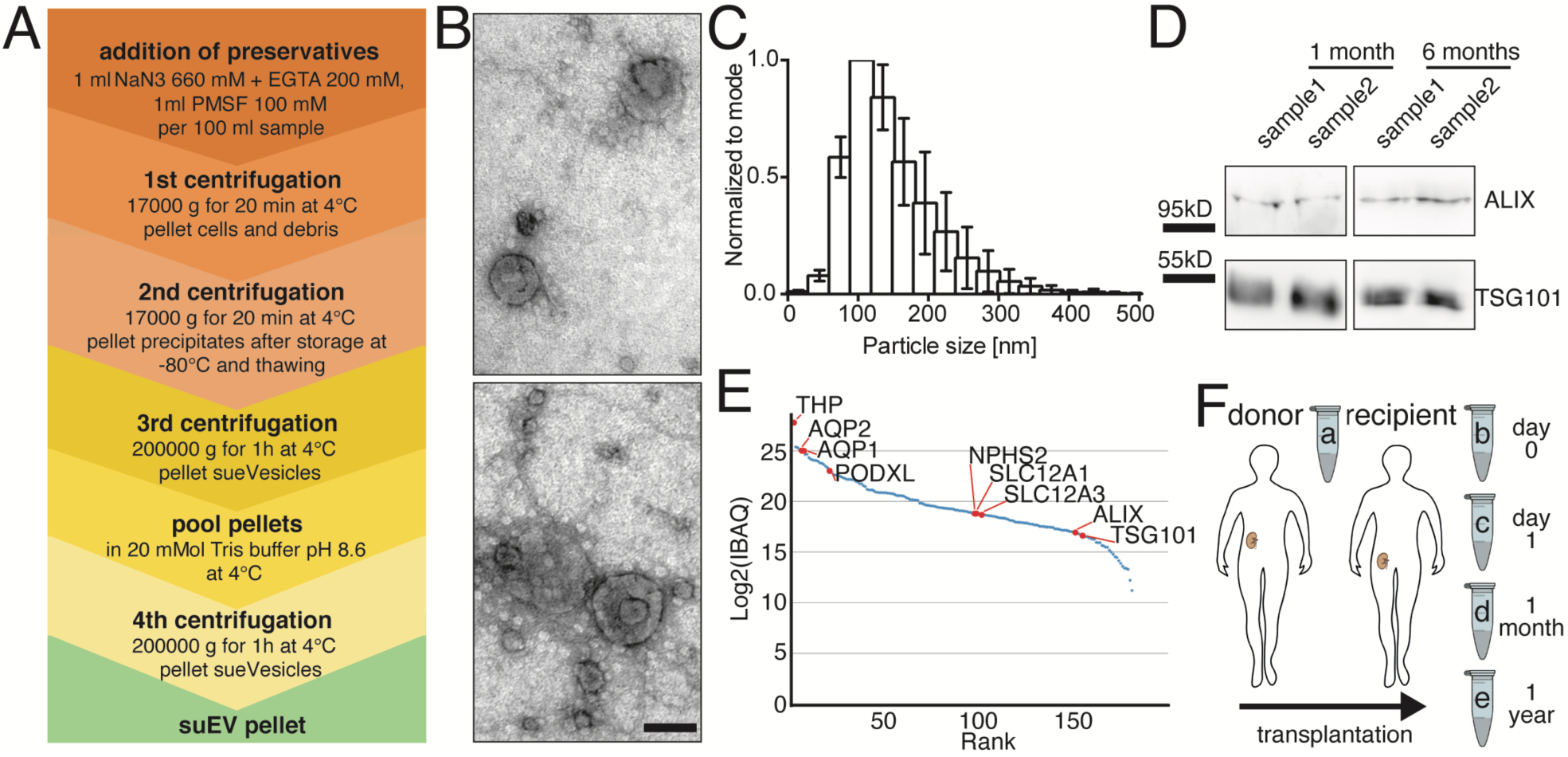
Differential centrifugation separates small urinary extracellular vesicles (suEVs) originating from all segments of the nephron. Schematic overview of the employed protocol of differential centrifugation **(A)**. Representative scanning electron microscopy of suEV pellets depicts EVs of typical exosomal cup shape and size and as well as smaller and bigger particles. Scalebar: 100nm **(B)**. Size distribution of suEVs measured in 4 independent suEV samples of healthy volunteers measured by Nanoparticle Tracking, bin: 30nm, errorbars: SD, **(C)**. Representative western blot analysis for exosomal markers ALIX and TSG101 in two separate suEV pellets after 1 and 6 months of storage in 8M Urea Buffer at −80°C **(D)**. Mass spectrometry analysis suEV pellet is able to marker proteins of all nephron segments. IBAQ: Intensity based absolute quantification (a.u.) **(E)**

### Clinical characteristics of donors and recipients of 22 living donor kidney transplantations (screening cohort)

We gathered suEV pellets over the course of 22 living donor transplantations. Figure 1F depicts a brief overview of the study. Urine was collected the day before transplantation from allograft donors (sample A) and their corresponding recipients directly after transplantation (sample B), 1 day after transplantation (sample C), 4 weeks (sample D) and one year after transplantation (sample E). Table 1 gives an overview of the clinical characteristics of the donors (Tbl. 1A) and corresponding recipients (Tbl. 1B). Medium age of donors was 51.5 years with roughly two thirds of the first cohort being women. Most donors showed a measured creatinine clearance of more than 100 ml/min/1,73m^2^ and only 3 donors suffered from arterial hypertension. Recipients were younger in age, mostly male and showed a lower body weight than the donors on average. More than two thirds underwent preemptive transplantation with the rest having been on dialysis between 2 and 39 months. Underlying renal diseases were almost tripartite between genetic, autoimmune and other causes with only one patient suffering from diabetic nephropathy. 72% of patients received basiliximab as an induction therapy and prednisolone, mycophenolic acid and tacrolimus for maintenance after transplantation. Three of 22 patients did not show a decrease in serum creatinine values of >1 mg/dl within the first 48 h and did not drop below a value of 3 mg/dl within the first week. For this study, these patients were classified as exhibiting delayed graft function for the following correlations.

**Table 1.** Patient characteristics (Cohort 1). Donors (**A**), Recipients (**B**)

| Category | Subcategory | Value | ± SD |
| --- | --- | --- | --- |
| **A (Donors)** |  |  |  |
| Age (in years) |  | 51,50 | ±11,09 |
| Sex |  |  |  |
|  | female (%) | 63,64 |  |
|  | male (%) | 36,36 |  |
| Weight (kg) |  | 76,95 | ±13,52 |
| measured clearance (ml/min/1,73m<sup>2</sup>) |  | 126,10 | ±32,21 |
| estimated GFR (CKD-EPI 2009) |  | 92,91 | ±12,64 |
| Hypertension (%) |  | 13,64 |  |
|  | ACE Inhibitors / AT1R Blockers (%)? | 13,64 |  |
| **B (Recipients)** |  |  |  |
| Age (in years) |  | 45,68 | ±15,86 |
| Sex |  |  |  |
|  | female (%) | 45,45 |  |
|  | male (%) | 54,55 |  |
| Weight (kg) |  | 69,89 | ±17,03 |
| Diuresis prior to transplant (%) |  | 68,18 |  |
| Dialysis prior to transplant (%) |  | 31,82 |  |
|  | time on dialysis in months | 15,29 | ±12,37 |
| Underlying renal disease |  |  |  |
|  | genetic (%) | 31,82 |  |
|  | diabetic (%) | 4,55 |  |
|  | glomerulonephritis (%) | 31,82 |  |
|  | other (%) | 31,82 |  |
| Retransplant (yes (number) / no) |  |  |  |
|  | yes (2nd transplant) (%) | 9,09 |  |
|  | no (%) | 90,91 |  |
| CMV status (rec/don) |  |  |  |
|  | neg/neg (%) | 18,18 |  |
|  | neg/pos (%) | 18,18 |  |
|  | pos/neg (%) | 13,64 |  |
|  | pos/pos (%) | 50,00 |  |
| Induction |  |  |  |
|  | ATG (%) | 9,09 |  |
|  | Basiliximab (%) | 90,91 |  |
| Initial Immunosuppression |  |  |  |
|  | Prednisolone / Cyclosporine A / MMF (%) | 27,27 |  |
|  | Prednisolone / Tacrolimus / MMF (%) | 72,73 |  |
| Cold ischemia time (in min) |  | 174,00 | ±26,80 |
| GFR after 6 months |  | 49,82 | ±15,58 |
| GFR after 12 months |  | 53,23 | ±16,88 |

### Unbiased proteomic analysis of first 22 transplant sets reveals temporal changes of the suEV proteome

Using mass spectrometry, we detected > 1700 individual proteins present in ≥ 50% of samples. The obtained data set is freely available at PRIDE/ProteomExchange (http://www.ebi.ac.uk/pride (*20*) – accession number: PXD005219). Hierarchical clustering revealed a strong tendency of samples collected at the same timepoint during transplantation to cluster together (Fig. 2A). Specifically, the two samples obtained early after transplantation (B and C) share a subcluster clearly distinguishing them from A, D and E samples – i.e. with the proteomic profiles one month and one year after transplantation showing more resemblance to the profile in the donor sample. In order to facilitate fast and uncomplicated access to a primary analysis and visualization of the data set, we have designed a browser-based application allowing the direct search for protein LFQs, their rank within the data set as well as the temporal changes in their abundance in suEVs during living donor kidney transplantation (http://shiny.cecad.uni-koeln.de:3838/suEV/).

**Fig.2:**
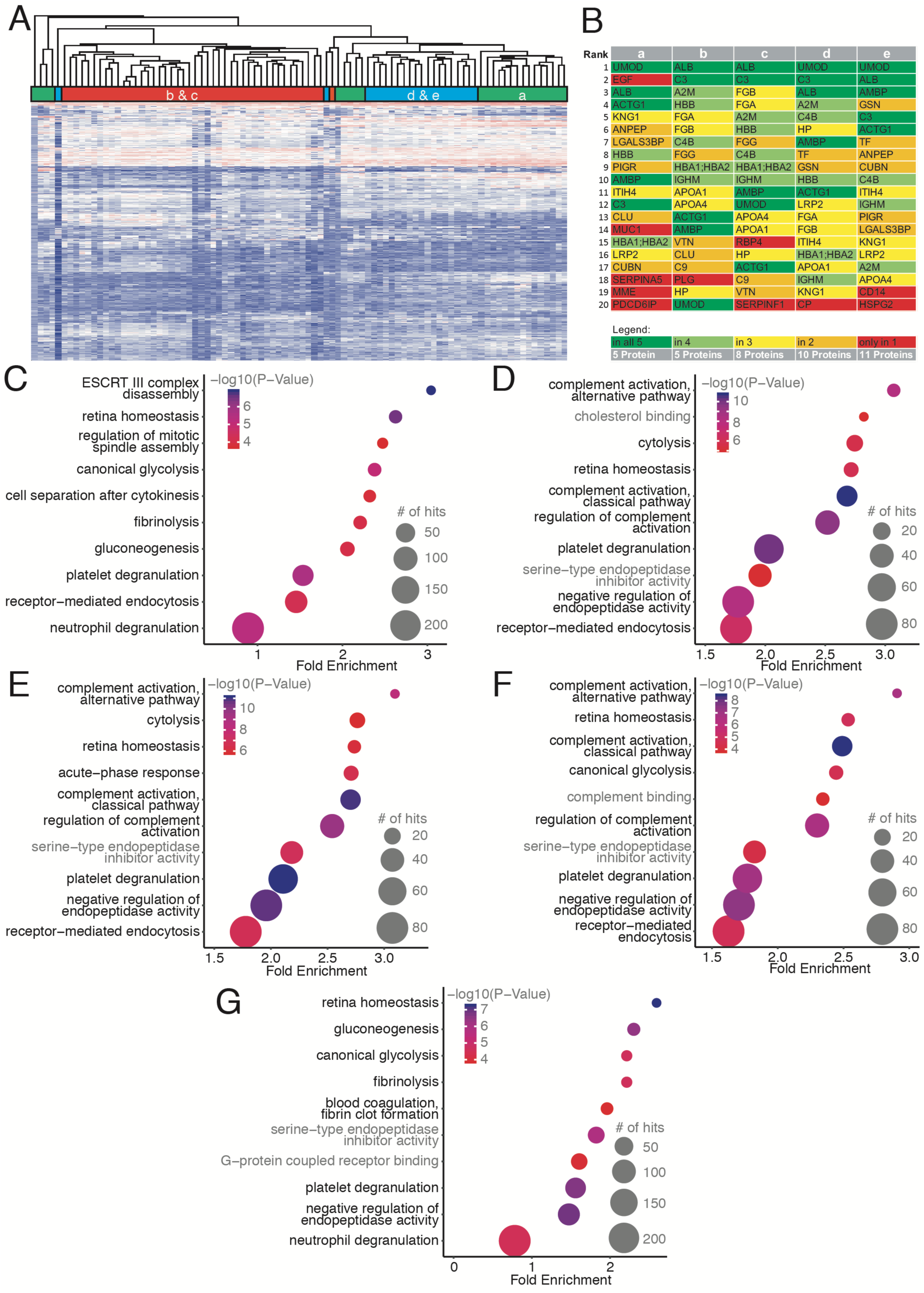
The suEV proteome clusters depending on the collection timepoint throughout living donor kidney transplantation. Hierarchical clustering of all single sample protein measurements of the first cohort **(A)**. Top 20 annotated proteins for each timepoint of sample collection **(B)**. Bubble plots of top 10 significant GO terms for each timepoint of sample collection. P value depicted as color code, number of annotated proteins corresponding to bubble size, black GO terms indicating biological processes, gray GO terms indicating molecular function. **(C:** Timepoint A – donor sample, **D:** Timepoint B – day 0, **E:** Timepoint C – day 1, **F:** Timepoint D – 1 month, **G:** Timepoint E – 1 year).

When comparing the top 20 proteins per timepoint ranked by label-free quantification after excluding proteins that could not be annotated, we detected 25% of these proteins to be present among the top 20 at all timepoints (Fig. 2B). These proteins contain the ubiquitous urinary protein uromodulin (UMOD) as well as albumin (ALB), AMBP and Complement Factor 3 (C3) but, interestingly, also cytoplasmatic actin (ACTG). Despite the hierarchical clustering results, the donor samples differ most from all other timepoints regarding the top 20 suEV proteins indicating a shift induced by transplantation itself. Programmed cell death 6-interacting protein (PDCD6IP), a bonafide marker for small extracellular vesicles, for instance, was only detected among the top 20 proteins in the donor samples. All early timepoints up to 4 weeks after transplantation (B/C/D) show an overrepresentation of proteins typically expected in serum rather than urine including fibrinogen chains alpha and beta and apolipoprotein A1 and A4.

To further dissect the impact of renal transplantation on the suEV proteome, we performed Gene Ontology (GO) analyses of all detected proteins per timepoint (Fig. 2C-G). We observed a striking overlap among the GO terms of highest significance per timepoint. “Platelet degranulation” and “Retina homeostasis” was detected as highly significant at all timepoints while “Receptor-mediated endocytosis” was absent, solely, at timepoint E (Fig. 2G). “Gluconeogenesis” was not annotated at timepoints b and c and not among the top 20 terms of highest significance at timepoint d while “Cytolysis” showed the exact opposite pattern. GO terms for complement activation and regulation were detected among the top 10 significant GO terms exclusively after transplantation (Fig. 2D, E & F) but lacking in the donor sample and decreased in significance 12 months after transplantation (Fig. 2C & G and http://shiny.cecad.uni-koeln.de:3838/suEV/).

We also observed a considerable overlap when ranking the top GO terms by enrichment (Fig. S2). However, proteins associated with cholesterol uptake and lipoprotein binding were only enriched in the samples early after transplantation (Fig. S2B & C) whilst proteins involved in Toll-like-receptor 4 and arachidonic acid binding were overrepresented in the donor sample and one year after transplantation (Fig. S2A & E).

In contrast to the GO analysis performed separately for each timepoint, GO terms for the difference in the suEV proteome content of each timepoint after renal transplantation compared to the initial donor suEV proteome showed much less of an overlap both ranked by enrichment and significance (Fig. S3/S4). Interestingly, we detected a significant fold change for proteins associated with the regulation of complement activation one day and one month after transplantation (Fig. S3B, C and S4A). Four weeks after transplantation, “cytolysis”, “clearance of apoptotic cells” and protein as well as membrane trafficking pathways were overrepresented (Fig. S4C).

To further elucidate the timepoint-dependent shift in the suEV proteome during renal transplantation, we analyzed the data set for clusters of proteins showing the same temporal signatures. We were able to identify 9 clusters exhibiting specific temporal shifts in their protein abundances (Fig. 3).

**Fig.3:**
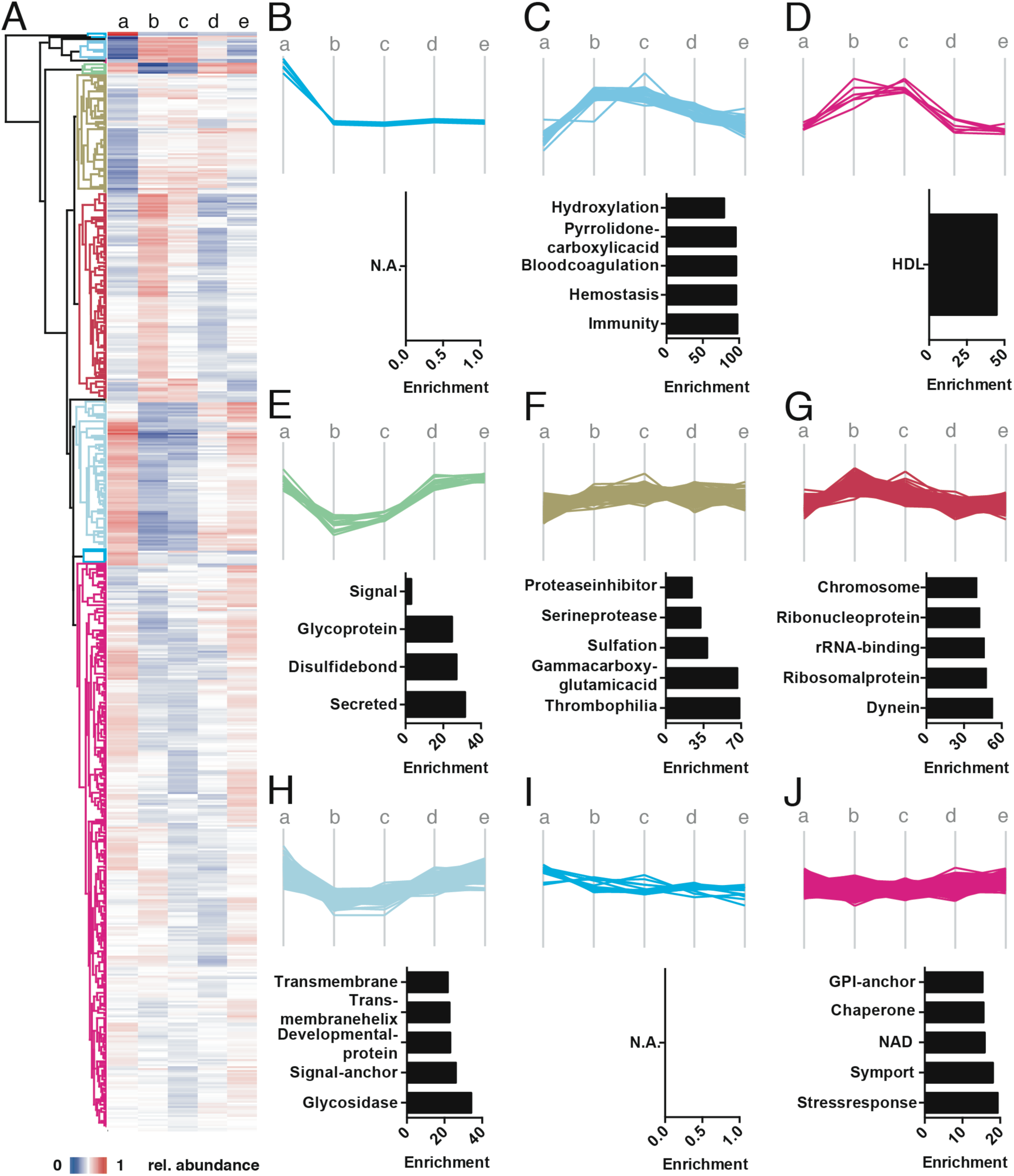
The suEV proteome underlies specific temporal changes indicative of specific biological processes in allograft adaptation. Hierarchical clustering of mean relative protein abundance per timepoint **(A)**. Spaghetti plots depicting suEV protein clusters with specific temporal changes throughout renal transplantation with enriched GO keywords bargraphs below each cluster. Color code indicates cluster branches on the heatmap depicted in 3A **(B-J)**.

Proteins with rising abundances after transplantation and a following decrease to levels above those detected in the initial donor suEV pellet appear to play a role in Immunity and Blood coagulation (Fig. 3C) while HDL-associated proteins decreased to donor values one year after transplantation. Proteins with a persistent increase after transplantation (Fig. 3F) were enriched in GO terms of protease homeostasis and “Thrombophilia”. An S shaped temporal signature (rise in abundance at timepoints B and C, decrease at timepoint D and normalization at timepoint E) was detected for proteins involved in ribosomal and chromosomal biology as well as “Dynein” activity (Fig.3G). Conversely, decreased abundances right after transplantation were observed for proteins enriched in GO terms for “Glycoprotein” and Secreted proteins (Fig.3E). These terms showed a “recovery” to donor values one year after transplantation, while the majority of proteins in the terms “Transmembrane”, “Glycosidase” and “Signal-Anchor” remained slightly decreased in the suEV pellet after one year (Fig. 3H). Interestingly, stress response proteins were detected within the cluster exhibiting the lowest to non-detectable changes in the suEV proteome (Fig. 3J). Two clusters, one with a pronounced and permanent decrease in abundance right after transplantation and one with a slow continuous decrease in abundance, could not be annotated using standard GO terms (Fig. 3B and I).

### Identification of potential biomarkers predicting kidney function 6 and 12 months after renal transplantation using a leave-one-out cross validation model

To assess the potential of the acquired proteomics data for identifying novel biomarkers of the outcome of renal transplantation, we performed a correlative analysis with the estimated glomerular filtration rate (eGFR) 6 months and 12 months after renal transplantation using a leave-one-out cross validation model (for a description of the approach see Supplementary movie S1). We investigated the predictive value of the foldchange of protein abundances between timepoints A to B and A to C as well as the single protein intensities at timepoints A and C normalized to the mean protein intensity. The resulting candidate protein lists were filtered for I) the 10 candidates exhibiting the lowest error (lowest RSS squared) and highest stability (lowest stdev RSS squared) when correlated to the estimated GFR after 12 month and, II) the 20 candidates exhibiting the lowest error correlating both with the estimated GFR 6 and 12 months after transplantation (Supplementary Table S1).

To generate a candidate list of markers for further validation we complemented this selection by 5 proteins specifically not detected and 10 proteins with significantly higher expression in patients with delayed graft function at timepoints A and C (Table S1). These proteins, the 10 candidates described under I) and the overlap of proteins found both predictive for the eGFR after 6 and 12 months described under II) resulted in a target list of 64 proteins (Tbl. 2).

**Table 2.** Candidate Biomarker Protein List.

| List origin | Protein names |
| --- | --- |
| **Correlating with 12 Month GFR with lowest RSSsquare mean and lowest SD** | Galactokinase |
|  | BH3-interacting domain death agonist;BH3-interacting domain death agonist p15;BH3-interacting domain death agonist p13;BH3-interacting domain death agonist p11 |
|  | Pancreatic secretory granule membrane major glycoprotein GP2 |
|  | Prominin-1 |
|  | V-type proton ATPase subunit S1 |
|  | Myocilin |
|  | Eukaryotic translation initiation factor 3 subunit C;Eukaryotic translation initiation factor 3 subunit C-like protein |
|  | 40S ribosomal protein SA |
|  | Complement C1q subcomponent subunit B |
|  | Tetranectin |
|  | 2-deoxynucleoside 5-phosphate N-hydrolase 1 |
|  | Charged multivesicular body protein 2a |
|  | DnaJ homolog subfamily A member 2 |
|  | Cadherin-16 |
|  | Monocyte differentiation antigen CD14;Monocyte differentiation antigen CD14, urinary form;Monocyte differentiation antigen CD14, membrane-bound form;Monocyte differentiation antigen CD14 |
|  | Collagen alpha-2(IV) chain;Canstatin |
|  | Glutathione S-transferase A2;Glutathione S-transferase |
|  | Villin-1 |
|  | Excitatory amino acid transporter 3 |
|  | 26S protease regulatory subunit 8 |
|  | Spectrin beta chain, non-erythrocytic 1 |
|  | Galectin-3-binding protein |
|  | Receptor-type tyrosine-protein phosphatase eta |
|  | Importin subunit beta-1 |
|  | Protein phosphatase 1 regulatory subunit 7 |
|  | Phosphoenolpyruvate carboxykinase [GTP], mitochondrial |
|  | Decorin |
|  | Ragulator complex protein LAMTOR1 |
|  | Annexin;Annexin A4 |
|  | Alpha-(1,3)-fucosyltransferase |
|  | Protocadherin Fat 4 |
|  | Sortilin-related receptor |
|  | SLIT and NTRK-like protein 1 |
|  | Protein amnionless |
|  | Complement C1r subcomponent-like protein |
|  | Peflin |
| **Correlating with both 6 Month and 12 Month GFR with lowest RSSsquare** | IgGFc-binding protein |
|  | ATP-dependent RNA helicase DDX3X;ATP-dependent RNA helicase DDX3Y |
|  | Cartilage oligomeric matrix protein |
|  | BH3-interacting domain death agonist;BH3-interacting domain death agonist p15;BH3-interacting domain death agonist p13;BH3-interacting domain death agonist p11 |
|  | DnaJ homolog subfamily A member 2 |
|  | Phosphoenolpyruvate carboxykinase [GTP], mitochondrial |
|  | Haptoglobin-related protein |
|  | Eukaryotic translation initiation factor 3 subunit C;Eukaryotic translation initiation factor 3 subunit C-like protein |
|  | 40S ribosomal protein SA |
|  | Syntenin-1 |
|  | Fructosamine-3-kinase |
| **Top5-not detectable in delayed graft function** | Transcription elongation factor B polypeptide 1 |
|  | Ras-related protein Rab-27B |
|  | Sodium-dependent phosphate transport protein 2A |
|  | Calbindin |
|  | Copine-8 |
| **Top10-higher abundance in non delayed graft function** | Glutamate carboxypeptidase 2 |
|  | CD177 antigen |
|  | Protein-glutamine gamma-glutamyltransferase 4 |
|  | Arylsulfatase F |
|  | Semenogelin-1;Alpha-inhibin-92;Alpha-inhibin-31;Seminal basic protein |
|  | 40S ribosomal protein S17-like;40S ribosomal protein S17 |
|  | 60S ribosomal protein L23a |
|  | Histone H2B type 1-D;Histone H2B |
|  | Complement component C6 |
|  | Podocalyxin |
| **Extracellular vesicle markers** | TSG101 |
|  | ALIX |
|  | CD81 |
|  | CD40, Tumor necrosis factor receptor superfamily member 5 |
|  | Annexin A5 |

### Targeted mass spectrometry in suEVs of an independent validation cohort identifies PCK2 as a predictor of long-term transplant function

To check validity of these 64 suEV candidate markers we performed targeted mass spectrometry assays for these proteins. These assays were then used to measure suEV pellets collected in an independent validation cohort throughout 22 additional living donor transplantations. Table 3 gives an overview of the patient characteristics of this validation cohort.

**Table 3.** Patient characteristics (Cohort 2). Donors **(A)**, Recipients **(B)**

| Category | Subcategory | Value | ± SD |
| --- | --- | --- | --- |
| **A (Donors)** |  |  |  |
| Age (in years) |  | 53,77 | ±10,7 |
| Sex |  |  |  |
|  | female (%) | 50,00 |  |
|  | male (%) | 50,00 |  |
| Weight (kg) |  | 79,53 | ±14,27 |
| measured clearance (ml/min/1,73m<sup>2</sup>) |  | 68,90 | ±18,64 |
| estimated GFR (CKD-EPI 2009) |  | 55,41 | ±11,60 |
| Hypertension (%) |  | 31,82 |  |
|  | ACE Inhibitors / AT1R Blockers (%) | 31,82 |  |
| **B (Recipients)** |  |  |  |
| Age (in years) |  | 46,05 | ±15,06 |
| Sex |  |  |  |
|  | female (%) | 45,45 |  |
|  | male (%) | 54,55 |  |
| Weight (kg) |  | 81,21 | ±15,02 |
| Diuresis prior to transplant (%) |  | 77,27 |  |
| Dialysis prior to transplant (%) |  | 22,73 |  |
|  | time on dialysis in months | 16,24 | ±43,90 |
| Underlying renal disease |  |  |  |
|  | genetic (%) | 31,82 |  |
|  | diabetic (%) | 0,00 |  |
|  | glomerulonephritis (%) | 18,18 |  |
|  | other (%) | 50,00 |  |
| Retransplant (yes (number) / no) |  |  |  |
|  | yes (2nd transplant) (%) | 9,09 |  |
|  | no (%) | 90,91 |  |
| CMV status (rec/don) |  |  |  |
|  | neg/neg (%) | 18,18 |  |
|  | neg/pos (%) | 18,18 |  |
|  | pos/neg (%) | 13,64 |  |
|  | pos/pos (%) | 50,00 |  |
| Induction |  |  |  |
|  | ATG (%) | 0,00 |  |
|  | Basiliximab (%) | 100,00 |  |
| Initial Immunosuppression |  |  |  |
|  | Prednisolone / Cyclosporine A / MMF (%) | 22,73 |  |
|  | Prednisolone / Tacrolimus / MMF (%) | 72,73 |  |
| Cold ischemia time (in min) |  | 172,70 | ±23,62 |
| GFR after 6 months |  | 53,33 | ±16,18 |
| GFR after 12 months |  | 59,16 | ±20,50 |

Renal function as indicated by both measured creatinine clearance (68,9 ml/min/1,73m^2^) and estimated GFR (55,41 ml/min/1,73m^2^) was lower in comparison to the first cohort with 7 patients being hypertensive and receiving corresponding medication (Tbl. 3A). Recipients were heavier than those of the initial cohort but otherwise closely resembled the latter, also when examining the renal function 6 months and 12 months after transplantation (Tbl. 3B).

We again performed both our correlative analysis and filter algorithm to identify proteins with a robust correlation to renal function 6 and 12 months after transplantation (supplementary table S2). This validation experiment lead to the confirmation of phosphoenolpyruvate carboxykinase 2 (PCK2 / PCK2) as an early marker predicting transplant outcome after 1 year. PCK2 abundance in suEVs 1 day after transplantation (timepoint C) showed a robust positive correlation with the estimated GFR 12 months after renal transplantation in both cohorts (Fig. 4A & B).

**Fig.4:**
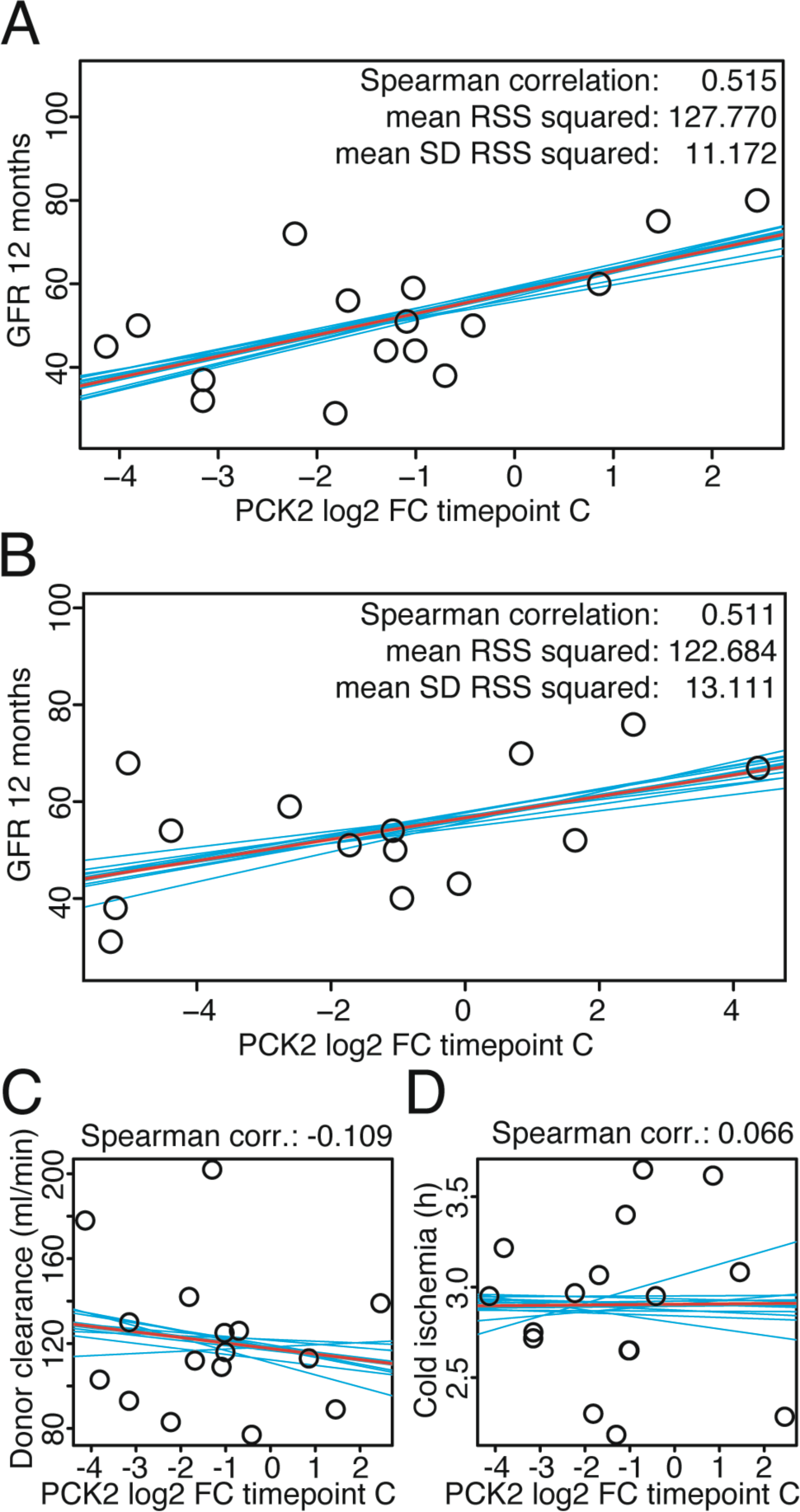
PCK2 abundance in suEVs on day one after transplantation correlates with estimated GFR 12 months after transplantation. Correlation plots of PCK2 intensity foldchange to mean intensity at timepoint C to GFR 12 months after transplantation measured in the initial **(A)** and validation **(B)** transplant cohort. Correlation plots of PCK2 intensity foldchange to mean intensity at timepoint C to donor clearance **(C)** and cold ischemia **(D)** measured in the initial cohort. Blue: individual linear regression models; Red: merged linear regression model for all samples.

We detected no predictive value for PCK2 abundances in suEVs of the donor sample or the foldchanges directly after and 1 day after transplantation (Fig. S5A-E). Strikingly, the abundance of PCK2 in suEVs one day after transplantation did not correlate with either the initially measured GFR of donors (Fig. 4C) or the recorded cold ischemia time (Fig. 4D) showing its potential as an independent indicator.

### suEV abundance of PCK2 rises initially during renal transplantation but does not reflect renal tissue PCK2 levels

In the initial unbiased proteomic measurement, PCK2 was detected at rank 1515 of 1728 with a mean LFQ of 21.88 at timepoint A (see (http://shiny.cecad.uni-koeln.de:3838/suEV/). Consequently, it appeared important to prove its localization within suEVs. Using immunogold labelling and transmission electron microscopy, we detected PCK2 within a subset of suEVs of 100-400nm diameter (Fig. 5A). Vesicles were permeabilized for 30 minutes (see materials and methods) allowing antibody penetration into the vesicular lumen for successful staining. No staining was detected within non-permeabilized vesicles (Fig. S6A).

**Fig.5:**
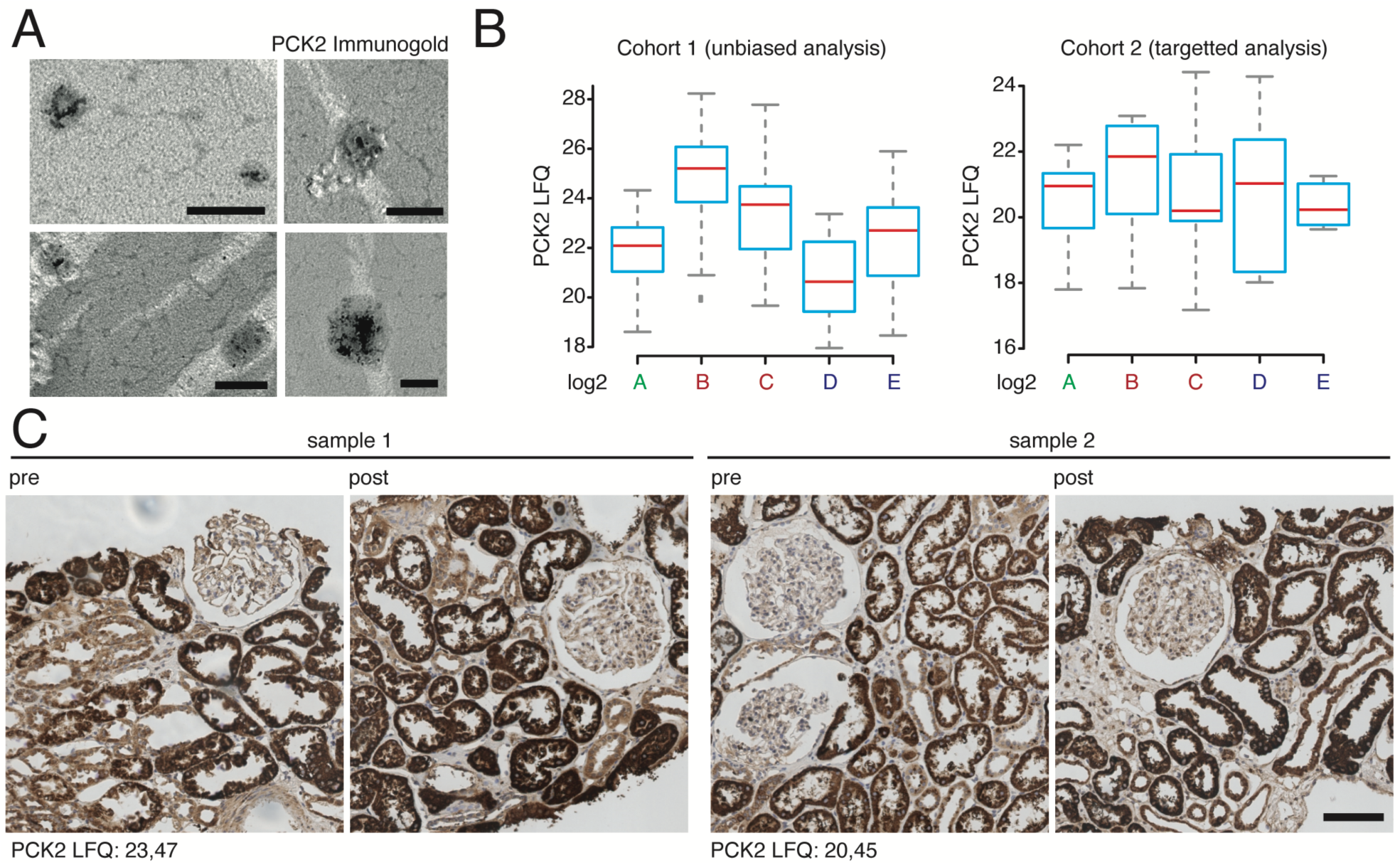
PCK2 localizes to suEVs, increases during the initial stages of transplantation and does not correlate to tissue PCK2 levels after reperfusion. Electron microscopy depicting permeabilized small urinary extracellular vesicles harbouring immunogold labelled PCK2, scalebar 200nm **(A)**. Box plot diagrams depicting suEV PCK2 abundance at different timepoints throughout living donor kidney transplantation in the initial and validation cohort. Mean (red line), 95% confidence interval (blue box) **(B)**. Immunohistochemistry staining for PCK2 in renal biopsies taken from two donors / patients (sample 1 / 2) included in the suEV analysis before explantation (pre) and after reperfusion (post). suEV PCK2 LFQs at timepoint C indicated in the lower left corner, scalebar 100µm **(C)**.

PCK2 shows a rising abundance in suEVs until 4 weeks after transplantation, with a decrease towards values measured in the donor sample one year after transplantation (Fig. 5B). To examine, whether the changes in suEV levels of PCK2 were due to differences in renal expression, we performed immunohistochemistry for PCK2 in renal biopsies taken during living donor kidney transplantation before explantation (pre) and right after reperfusion (post) from a subset of patients also included in the suEV analysis. We detected a strong signal predominantly in proximal tubules, as described before, but no overt changes after reperfusion or evidence for a correlation to the measured LFQs in the suEV proteome of the donor samples (Fig. 5C).

## Discussion

We present the first quantitative analysis of the temporal changes in the suEV proteome throughout living donor kidney transplantation. Despite the acceptance of kidney transplantation as the preferred renal replacement therapy in ESRD in developed countries, the precise effects cold ischemia time and reperfusion exert on renal cells are incompletely understood. Mouse models can only partially shed light on these mechanisms, as their organism differs greatly concerning their immune systems, one of the key players involved in transplant biology (*21*).

Nevertheless, a better understanding could impact future therapeutic approaches to renal transplantation and lead to a better overall outcome both of organ function in the early days and long-term organ survival. This study represents the first in-human analysis of these proteomic patterns and allows novel insights into the molecular processes taking place in the early days and weeks after transplantation as well as those underlying the long-term recovery. Former studies in donors of renal allografts investigated the correlation between EVs carrying specific markers to the tissue characteristics obtained through renal biopsy during transplantation(*22*). While an impressive number of patients was investigated and valuable information was obtained, a deeper understanding was limited due to the use of a specific antibody panel.

Whole urine proteomics have been employed in the setting of renal transplantation in the past (*23–25*). Yet, it is important to note, that, apart from a “core urinary proteome”, free urinary proteins showed a fairly high inter-and intra-individual variability (*26*). Employing an unbiased proteomic analysis, we were able to detect >1700 proteins enriched through our separation protocol.

The suEV proteome showed a strong tendency to form timepoint-dependent subclusters. This finding confirms technical reproducibility in a clinical setting, as the individual vesicle separation was performed by various trained members of our group, and the storage time of the initially collected urine samples varied from sample to sample.

When considering the top 20 proteins of highest abundance, 5 proteins were detectable to a high degree at all timepoints. While UMOD, ALB, C3 could be interpreted as a sign for co-precipitation of free proteins, it is worth noting that all three proteins were also present in proteomic analyses of urinary extracellular vesicles separated through protocols involving a sucrose cushion or immune affinity (*18, 27, 28*). Interestingly, PDCD6IP or ALIX, which, as a bona fide marker for the late endosome and the vesicles derived from it, is an integral interactor with the ESCRT machinery in vesicle formation, was only present among the proteins of highest abundance in suEVs before transplantation. This also holds true when broadening the filter to the top 50 proteins, indicating a decrease in ALIX-positive vesicles with potential implications for vesicle formation and late endosome maturation throughout renal transplantation. Further indication for an alteration in the release of vesicles is given by the reduction in TSG101 right after transplantation as TSG was proposed as a surrogate marker for urinary vesicle number in a recent study (*29*).

The surge in serum-associated proteins at timepoints B-D can reflect differences in EV packing as well as a decreased intercellular barrier upon reperfusion resulting in the crossing of plasma extracellular vesicles into the urinary space (*30, 31*). Yet, the fact that these proteins are still detected as late as 4 weeks after transplantation (timepoint d), speaks for them rather to be derived from cells of the nephron. Recently, the channel proteins AQP1 and AQP2 were shown to be decreased in urinary EVs after ischemia reperfusion injury in a rat model (*32*) and in human patients undergoing renal transplantation (*33*). We were able to reproduce these findings in our unbiased proteomic approach.

Regarding the GO term analyses, the term “platelet degranulation” was detected as highly significant at all timepoints (Fig. 2C-G). As the common denominator of most proteins associated to the term is their incorporation in platelet-derived extracellular vesicles (*31*), we believe that there is constant passage of these vesicles into the urinary space either through the glomerular filtration barrier, leaky intercellular space or active transcellular transport.

The surge in proteins associated with both complement activation, its regulation and cytolysis most likely reflects the initial immune response triggered by both ischemia-reperfusion and exposure to the allograft. This allows for the hypothesis that an early blockade of these responses – probably focusing on the complement system – may be beneficial (*34, 35*). Furthermore, the analysis of suEVs could become a diagnostic strategy to monitor complement activation in the setting of renal transplantation.

Toll-like receptor 4 signaling has long been investigated as a mediator for renal fibrosis after transplantation and over the course of various nephropathies (*36–41*). In contrast to that, derivatives of arachidonic acids and their mimics were shown to exhibit anti-fibrotic effects and polymorphisms in their CYP-dependent metabolism to influence susceptibility to renal injury (*42–44*).

GO term analysis of the differences in the suEV proteome of each timepoint to the healthy (donor) state revealed the vast and quickly adapting changes in EV assembly and cargo. Again, the significant enrichment of complement pathway-activating proteins is not surprising and confirms published data based on tissue analyses (*34*). Conversely, the enrichment of proteins associated to “multivesicular body assembly” and “viral (vesicle) budding” point to a heightened activity of EV production, potentially triggered through the ischemia reperfusion injury, which is also reflected by a continuous rise in TSG protein abundance after an initial dip right after transplantation ((http://shiny.cecad.uni-koeln.de:3838/suEV/). Induction of vesicle production has been described both in cell culture damage systems as well as cardiac ischemia in humans and mice and has been associated with the recruitment of monocytes to the damaged tissue (*45, 46*). Given the fact that EV administration has been shown to facilitate renal recovery after ischemia-reperfusion damage(*47*), our finding could point towards an intrinsic counter mechanism.

Besides being a resource for the biological temporal changes after transplantation, we hypothesized that the suEV proteome could be a future source for the identification of biomarkers. A set of proteins detected in EVs early after transplantation correlated clearly with eGFR 6 and 12 months later. Since the robust identification of marker proteins cannot be achieved using untargeted proteomics in a limited set of patients, we then aimed for the validation in a separate cohort using targeted mass spectrometry. Here, PCK2 abundance in the suEV pellet one day after transplantation was confirmed to correlate with eGFR one year after transplantation (Fig. 4A&B). This correlation was not detected when measured at any other timepoint or investigating the difference in abundance between timepoints (Fig. S5). Strikingly, PCK2 abundance did not correlate with the initial donor clearance or cold ischemia time during transplantation (Fig.4C&D).

This finding points towards unknown mechanisms in transplant adaptation or resistance to ischemia-reperfusion, revealed by the investigation of the suEV proteome. Mechanistically, PCK2 has been shown to react to states of metabolic starvation, particularly in cancer. In glucose deprivation, PCK2 upregulation with ensuing gluconeogenesis stabilizes nutrient supply and tumor growth with growth impairment upon PCK2 inhibition(*48, 49*). This process may well take place also in the renal allograft, as PCK2 is highly expressed in the proximal tubule and the tubular compartment is the prime target of ischemic and acute damage in the kidney (*50*).

An increase in PCK2 levels could potentially counteract the observed decrease in gluconeogenesis (Fig.2C-G) at timepoints B, C and D as indicated by our GO analysis. It is possible that this PCK2 induction is activated after reperfusion, but takes time to reach a measurable peak in tissue analyses. This would explain why we did not detect evidence for a correlation of tissue PCK2 levels right after reperfusion in immunohistochemistry with suEV PCK2 levels one day after transplantation. Albeit, there is ongoing debate as to whether suEV protein content does correlate with tissue protein content at all (*51, 52*). PCK2 is a mitochondrial protein. An alternative explanation of the increased PCK2 content in suEVs without a change in the cellular content may be an activation of mitochondrial activity in individuals protected from damage and a consecutive rise in the release of this protein. However, this remains speculation at this point and will require further cell biological studies.

The presented study entails limitations due to the separation protocol and the patient cohort chosen. Due to the protocol chosen, we cannot draw conclusions of the actual vesicle type contributing to the protein signals. To this day, there is no agreed gold standard separation protocol and most studies still use the widely employed protocol of differential (ultra-)centrifugation (*15*)(*13*). We, as well, chose a modification of these protocols, due to several reasons. Differential ultracentrifugation enables the use of higher volumes of the initial sample and, therefore, the separation of more EVs (*18*). Differential ultracentrifugation fails to separate a pure vesicle fraction, yet filtration and precipitation techniques were shown to unspecifically concentrate free urinary proteins as well while often excluding specific EV fractions from the separated pellet. More laborious techniques such as sucrose cushion gradient separation, would make a translation to the clinical setting extremely difficult.

The fact, that patients undergoing living donor transplantation were chosen, leads to a collective experienced with only short cold ischemia periods and very good outcome, that is not necessarily comparable in deceased donor transplantations. Nevertheless, this allowed us to include samples of donors with precisely monitored renal health and follow the effects of early and long-term transplant adaptation for up to one year.

Whilst our study shows the potential of suEVs as a source of biomarkers, the small number of patients (22 in the screening cohort, 22 in the validation cohort) does not allow for definite statement on the predictive capacity of suEV PCK2. Therefore, a potential use of this marker will depend on future larger trials. Our data set can serve as the first step to inform the design of such studies. Since PCK2 was located to suEVs between 100-400nm in size, future analyses can employ separation methods not relying on ultracentrifugation such as size exclusion chromatography to open the investigation of PCK2 in suEVs to a broad range of groups worldwide. Likewise, the timepoint of measurement opens up the possibility to validate the predictive value in deceased donor transplantation.

In conclusion, using differential (ultra-) centrifugation we assembled and analyzed a concise atlas reflecting the temporal changes in the proteome of small urinary extracellular vesicles throughout the course of living donor kidney transplantation. We detected specific temporal profiles, e.g. a surge in complement-associated proteins closely after transplantation reflective of early immunologic effects. Furthermore, our approach revealed the abundance of phosphoenol pyruvate carboxykinase (PCK2) in the suEV proteome one day after transplantation to have a predictive value for overall kidney function one year after transplantation. This study underlines the potential of analyzing suEVs for monitoring immune response activity and for biomarker discovery using a fast and easy protocol for their separation.

## Materials and Methods

### Patient recruitment

Patients were recruited through the Department of General, Visceral and Cancer Surgery, Division of Transplantation Surgery, Transplant Center Cologne according to the approved ethics statements 11-310 and 15-032, Ethics committee, University of Cologne, Cologne, Germany. The performed studies are registered at the German Clinical Trials Registry under the ID: DRKS00010534 and DRKS00007704.

### Urine collection and pre-processing

≥ 50ml urine samples were either collected as second morning midstream sample (timepoint A, D and E) or as catheter urine from a fresh collection bag (timepoint B and C) into standard urine sample cups (Sarstedt, Germany). Preservatives and protease inhibitors were added to the final concentrations 6,6mM NaN_3_, 2,0mM EGTA, 1,0mM PMSF within 90 minutes after sample collection. Afterwards cells, debris and large to medium sized EVs were separated via high speed centrifugation using a Beckman Avanti Centrifuge with a TLA 16,250 fixed angle rotor in 50ml centrifugation Tubes (Greiner Bio-One, Germany) at 17.000g, 4°C for 20 minutes. Supernatant was transferred to fresh 50ml centrifugation tubes and samples stored at −80°C until further processing.

### Small urinary extracellular vesicle separation

For suEV separation, frozen samples were thawed with subsequent repetition of centrifugation at 17.000g, 4°C for 20 minutes to pellet protein precipitates due to the freeze thaw cycle. Supernatant was afterwards transferred to polycarbonate ultracentrifugation bottles (Beckman Coulter, USA) and suEVs were separated at 200.000g, 4°C, 60 minutes using a Beckman Coulter Optima Ultracentrifuge with a 70Ti fixed angle rotor. Supernatant was discarded and suEV pellets resuspended and pooled in 4°C 20mM Tris Buffer pH 8,6 with subsequent repetition of the ultracentrifugation step. After discarding the supernatant, the suEV pellet of originally ≥25ml whole urine was resuspended in 50µl of Paraformaldehyde and stored at 4°C for electron microscopy or 40µl 8M Urea Buffer + Protease Inhibitor Mix (Sigma-Aldrich, USA) and stored at −80°C for protein analysis.

### Electron microscopy and assessment of size distribution

Samples were processed according to a modified version of previously published protocols (*53*). In short: suEV pellet was resuspended in 50µl 2% PFA (weight/vol) in PBS and incubated at 4°C over night. 5µl were added to formwar-coated grids and dried for 20 minutes at room temperature. Grids were washed 7 times in PBS for 2 minutes at room temperature before incubation in 1% glutharaldehyde (vol/vol) for 5 minutes. After rinsing in distilled water, grids were contrasted in 1,5% uranylacetate in distilled water (weight/vol) in the dark for 4 minutes and dried afterwards. Grids were analyzed at the TEM109 from Zeiss with a CCD camera by Tröndle. Size distribution was measured in 5 randomly picked images from two individual healthy subjects using the measuring tool in FIJI (*54, 55*).

### Electron microscopy and immunogold labelling of PCK2

For immunogold staining, 5 μl of resuspended paraformaldehyde fixed suEVs were transferred on Formvar-carbon coated nickel grids (20 minutes incubation). After washing with PBS the samples on the grids were permeabilized with 0.05% Tween (in the blocking solution) and blocked with 0.5% BSA for 30 minutes. suEVs were incubated with the primary rabbit-anti-PCK2 antibody (Abcam – ab187145) diluted 1:200 for 2 hours at room temperature, washed and labeled with a goat-anti-rabbit secondary antibody conjugated to 1.4 nm gold particles diluted 1:200 (Jackson ImmunoResearch - 711-205-152) for 1 hour at room temperature. After washing nanogold particles were enhanced for 2 minutes using HQ silver kit (Nanoprobes). Grids were contrasted as described above. Imaging was performed using a Philipps CM 100 TEM and Olympus imaging software ITEM.

### Western blot analysis

suEV pellets resuspened in 8M urea buffer were diluted using 1% triton X-100 buffer and 4% SDS sample buffer was added accordingly. Size separation was done using SDS-PAGE. Samples were blotted onto polyvinylidene difluoride membranes and visualized with enhanced chemiluminescence after incubation of blots with corresponding antibodies (anti-ALIX antibody BD Bioscience - 611620, anti-TSG101 antibody Abcam – ab30871).

### Unbiased proteomic analysis

Peptides were analyzed using a Q Exactive Plus tandem mass spectrometer (Thermo scientific). A 50 cm column was packed with 1.7 μm C18 beads (Dr Maisch GmbH). The column was used in a column oven. Peptides were fractionated by nano-liquid chromatography. For this purpose, a binary buffer system was used. This consisted of two buffers: buffer A: 0.1% formic acid and buffer B (80% acetonitrile, 0.1% formic acid). Linear gradients of 150 minutes duration were used according to the following scheme: 7-38% B, 80% B for 5 minutes and re-equilibration 5% B. Peptides were sprayed by Electron Spray Ionization (ESI) into a Q Exactive Plus Tandem mass spectrometer (*56*). The following parameters were used for MS1 scans: injection time 20 ms, resolution 70,000, mass range 200-1200 mz-1, 1E6 as to AGC target. MS2 scans were triggered by a top 10 method for further peptide fragmentation by means of higher-energy collisional dissociation (HCD) fragmentation. The following parameters were used for MS2 scans: 35,000 at 200 mz-1, AGC target to 5E5, max injection time 120 ms. Raw data (.raw) was analyzed using MaxQuant V 1.5.0.1 (*57*). The search (target decoy strategy) was performed against a 2017 reference human uniprot database. The default settings were used and the “MQLFQ” and “Match between runs” options were both used (*58*). The FDR cutoffs were FDR 0.01 for all peptide, PSM, and protein identifications. Perseus (v. 1.4.0.1 and v. 1.5.2.4) (*59*) was used for further analysis. To correct for multiple testing, a SAM approach was chosen (cutoff s0 = 1, FDR <0.001) (*60*). Significantly enriched GeneOntology terms were determined by Fisher’s Exact Test (FDR = 0.05).

### Correlative analysis

Detected proteins where correlated by linear regression models to eGFR. The eGFR was estimated using the CKD-EPI Creatinine Equation (*61*). Models were generated using a leave-one-out crossvalidation procedure(*62*). Here, all values but one are used as the training set (n-1) to calculate a linear model using the R function lm. The left out measurement (the “test set”) is calculated using method predict.lm and the difference between predicted and true value is used to calculate the prediction error of the model. The process is repeated according to the number of measured proteins (n) resulting in n linear models. Using these models a mean linear model is generated for each detected protein. From these models candidate proteins were determined for targeted mass spectrometry with criteria described in the main text. The targeted mass spectrometry of the independent validation cohort was analysed using the same approach.

### Targeted proteomic analysis

A parallel reaction monitoring PRM assay was set up. Analysis was performed using the LC-MS system as described for untargeted proteomics. Acquisition parameters for targeted proteomics data were as follows: 70 peptides were selected for scheduled PRM analysis (Supplementary Table S3). MS2 resolution was set at 140k. The AGC target was set at 1,000,000 at a maximum injection time of 100 ms. Peptides were isolated using an isolation window of 1.0 Th and fragmented with a normalized collision energy of 27. The two PCK2 peptides with the amino acid sequence HGVFVGSAMR (charge state 2, m/z = 530.771468, retention time = 31.92 min and EGALDLSGLR, charge state 2, m/z = 515.780013, retention time = 46.77 min were used. A scheduled method was composed using Skyline software environment (*63, 64*). A 4 min retention time window was used for acquisition on a 1h gradient that was previously described. Analysis and quantification of targeted proteomics data was performed using skyline. A library of 6 representative suEV rawfiles from the untargeted analysis was used as to construct the spectral library. Peaks were integrated and quantified using skyline. Normalized intensities were exported and processed. Only peptides within a 5 ppm mass window were considered.

### Immunohistochemistry

The tissue was fixed in 4% paraformaldehyde in phosphate-buffered saline (PBS) followed by paraffin embedding for histology. All steps were performed at room temperature (RT) using an autostainer 480S and protocol by Thermo Scientific. In brief, after deparaffinization with xylene and rehydration by graded series of ethanol the slides were washed with TRIS-NaCl with Tween (Medac-diagnostica, Germany). Peroxidase was blocked (Ultra Vision Hydrogen Peroxidase Block, Medac-diagnostica, Germany) and a pretreatment with citrate buffer at pH6 followed (PT modul Buffer 1, Thermo Scientific). The primary antibody rabbit-anti-PCK2 antibody (Abcam – ab187145) was diluted with antibody diluent (Medac-diagnostica, Germany) 1:1000 and developed using poly-HRP-anti mouse/rabbit IgG (Bright Vision, Medac-diagnostica, Germany) and DAB away kit (Biocare Medical, Germany). All reagents were used according to manufacturer’s protocols.

### Nanoparticle tracking analysis

EVs were analyzed by the ZetaView PMX-120 device (Particle Metrix, Germany). All samples were either diluted 1:1000 or 1:5000 in PBS to a final volume of 1 ml to achieve an ideal particle per frame value of 140 to 200. Every measurement, three times 11 cell positions have been analyzed after video capturing 30 frames, each position. Autofocus was set and all samples were measured with a camera sensitivity of 75, using a shutter value of 100 and a constant temperature of 25°C. Videos were analyzed by the ZetaView software 8.05.10 SP1 with a minimal area of 5, a maximal area of 1000 and a minimal brightness of 25. The number of completed tracks in NTA measurements was always greater than 1000 (*65*).

## Supporting information

Tbl S1

Tbl S2

Tbl S3

## Supplementary Materials

**Table S1.: Potential proteomic suEV biomarkers for renal outcome**

**Table S2.: Correlative analysis of potential proteomic suEV biomarkers detected in targeted proteomic analysis**

**Table S3.: Selected peptide sequences for targeted proteomic analysis**

**Movie S1: Schematic overview of leave-one-out crossvalidation procedure.** After plotting of all measured values per timepoint or fold change analysis, all values but one are used to create a correlative model. Subsequently, the missing value is predicted with the generated model, with the differences between predicted and true value indicating the prediction error and the correlative value of the model. The process is repeated according to the number of initial values creating correlative models of that number. All created models are summarized into an average model per protein. By using this mean model, the predictive value of protein abundance or fold change to GFR is calculated.

**Fig.S1: Differential centrifugation leads to a decrease Uromodulin to TSG ratio and separates small tetraspanin-containing vesicles.** Size distribution of suEVs measured in SEM of 2 independent samples of healthy volunteers **(A)**. Western Blot Analysis and Densitometry of Uromodulin and TSG101 in 2 independent samples of healthy volunteers **(B)**. Flow cytometric bead assay for tetraspanins CD9, CD63, CD81 and Phosphatidylserine (PS) of 4 independent samples of healthy volunteers **(C)**.

**Fig.S2: Enriched GO term analysis of absolute protein quantification per timepoint.** Bubble plots of top 10 GO terms enriched for each timepoint of sample collection. P value depicted as color code, number of annotated proteins corresponding to bubble size, black GO terms indicating biological processes, gray GO terms indicating molecular function **(A:** Timepoint A**, B:** Timepoint B**, C:** Timepoint C**, D:** Timepoint D**, E:** Timepoint E).

**Fig.S3: Significant GO term analysis of protein fold changes compared to initial donor sample.** Top 10 significant GO terms for each timepoint after transplantation compared to the initial donor samples. P value depicted as color code, number of annotated proteins corresponding to bubble size, black GO terms indicating biological processes, gray GO terms indicating molecular function **(A:** Timepoint B vs A**, B:** Timepoint C vs A**, C:** Timepoint D vs A**, D:** Timepoint E vs A).

**Fig.S4: Enriched GO term analysis of protein fold changes compared to initial donor sample.** Top 10 GO terms enriched for each timepoint after transplantation compared to the initial donor samples. P value depicted as color code, number of annotated proteins corresponding to bubble size, black GO terms indicating biological processes, gray GO terms indicating molecular function **(A:** Timepoint B vs A**, B:** Timepoint C vs A**, C:** Timepoint D vs A**, D:** Timepoint E vs A).

**Fig.S5: PCK2 abundance in suEVs at other timepoints does not correlate with estimated GFR 12 months after transplantation.** Correlation plots of PCK2 intensity foldchange to mean intensity at timepoint A **(A)** and B **(B)** and foldchanges between timepoints A vs. B **(C)**, A vs. C **(D)**, and B vs. C **(E)**, to GFR 12 months after transplantation measured in the validation cohort. Blue: individual linear regression models; Red: merged linear regression model for all samples.

**Fig.S6: PCK2 localizes to suEVs, increases during the initial stages of transplantation and does not correlate to tissue PCK2 levels after reperfusion.** Electron microscopy depicting non permeabilized small urinary extracellular vesicles after immunogold staining for PCK2, scalebar 100nm

## Acknowledgments

We acknowledge the excellent technical support of Ruth Herzog, Beatrix Martini, Valerie Oberüber, Serena Greco-Torres, Israa Kambar, Frederik Sand, Constantin Rill, Charlotte Dellmann, Martin Vitus, Ugur Keser, Ismini Halver, Dervla Reilly, Eva Schulze, Vincent Köntges, Bastian Trinsch, Philip Lützen, Nora Krasniqi, Babak Mochtarzadeh and the Imaging Facility at the CECAD, Cologne.

## Author contributions

The research was designed by F.B., M.R., A.B., D.S. and R.-U.M. F.B., M.R., K.B., C.K., I.P., D.Bu., D.Ba., R.W., M.S., H.G, O.K., V.G.P. and C.E.K. conducted the experiments and acquired the data. F.B., M.R., K.B., C.K., D.B., C.E.K. and R.-U.M. analyzed the data. F.B. and R.-U.M. wrote the manuscript. F.B., M.R., V.G.P., O.K., D.Ba., T.B.H., B.S., T.B., D.S., C.E.K. & R.-U.M. revised the final version. All authors approved the final version of the paper.

## Data and materials availability

The obtained data sets are freely available at PRIDE/ProteomExchange (http://www.ebi.ac.uk/pride (*20*) - Project name: Longitudinal study of the small urinary extracellular vesicle proteome during living donor kidney transplantation; Project accession: PXD005219; Reviewer account details: Username, Password: 3FR3kDp8. For direct and visual access see http://shiny.cecad.uni-koeln.de:3838/suEV/.

