## supplemental figures and information for "The proteome of small urinary extracellular vesicles after kidney transplantation as an indicator of renal cellular biology and a source for markers predicting outcome"

### Supplementary Materials

**Table S1.: Potential proteomic suEV biomarkers for renal outcome**

**Table S2.: Correlative analysis of potential proteomic suEV biomarkers detected in targeted proteomic analysis**

**Table S3.: Selected peptide sequences for targeted proteomic analysis**

**Movie S1: Schematic overview of leave-one-out crossvalidation procedure**. After plotting of all measured values per timepoint or fold change analysis, all values but one are used to create a correlative model. Subsequently, the missing value is predicted with the generated model, with the differences between predicted and true value indicating the prediction error and the correlative value of the model. The process is repeated according to the number of initial values creating correlative models of that number. All created models are summarized into an average model per protein. By using this mean model, the predictive value of protein abundance or fold change to GFR is calculated.

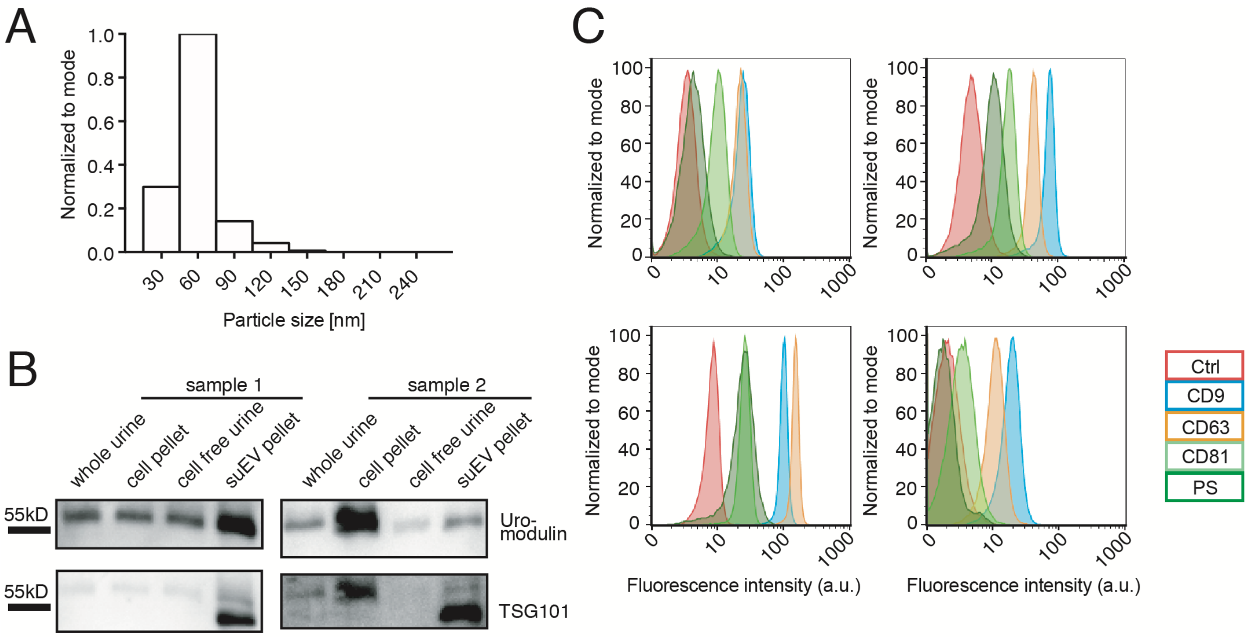


**Fig.S1: Differential centrifugation leads to a decrease Uromodulin to TSG ratio and separates small tetraspanin-containing vesicles.** Size distribution of suEVs measured in SEM of 2 independent samples of healthy volunteers **(A)**. Western Blot Analysis and Densitometry of Uromodulin and TSG101 in 2 independent samples of healthy volunteers **(B)**. Flow cytometric bead assay for tetraspanins CD9, CD63, CD81 and Phosphatidylserine (PS) of 4 independent samples of healthy volunteers **(C)**.

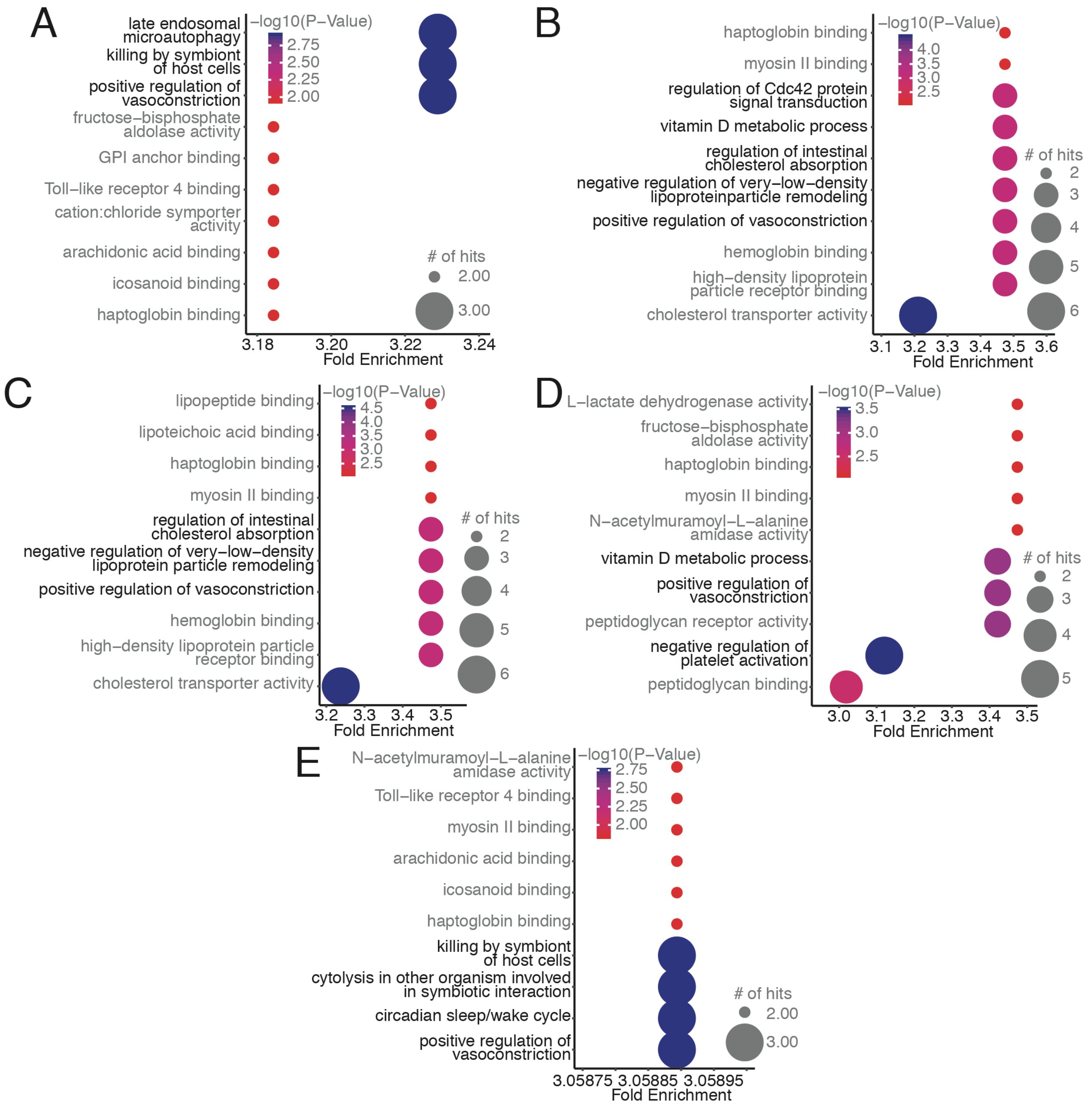


**Fig.S2: Enriched GO term analysis of absolute protein quantification per timepoint.** Bubble plots of top 10 GO terms enriched for each timepoint of sample collection. P value depicted as color code, number of annotated proteins corresponding to bubble size, black GO terms indicating biological processes, gray GO terms indicating molecular function **(A:** Timepoint A**, B:** Timepoint B**, C:** Timepoint C**, D:** Timepoint D**, E:** Timepoint E**)**.

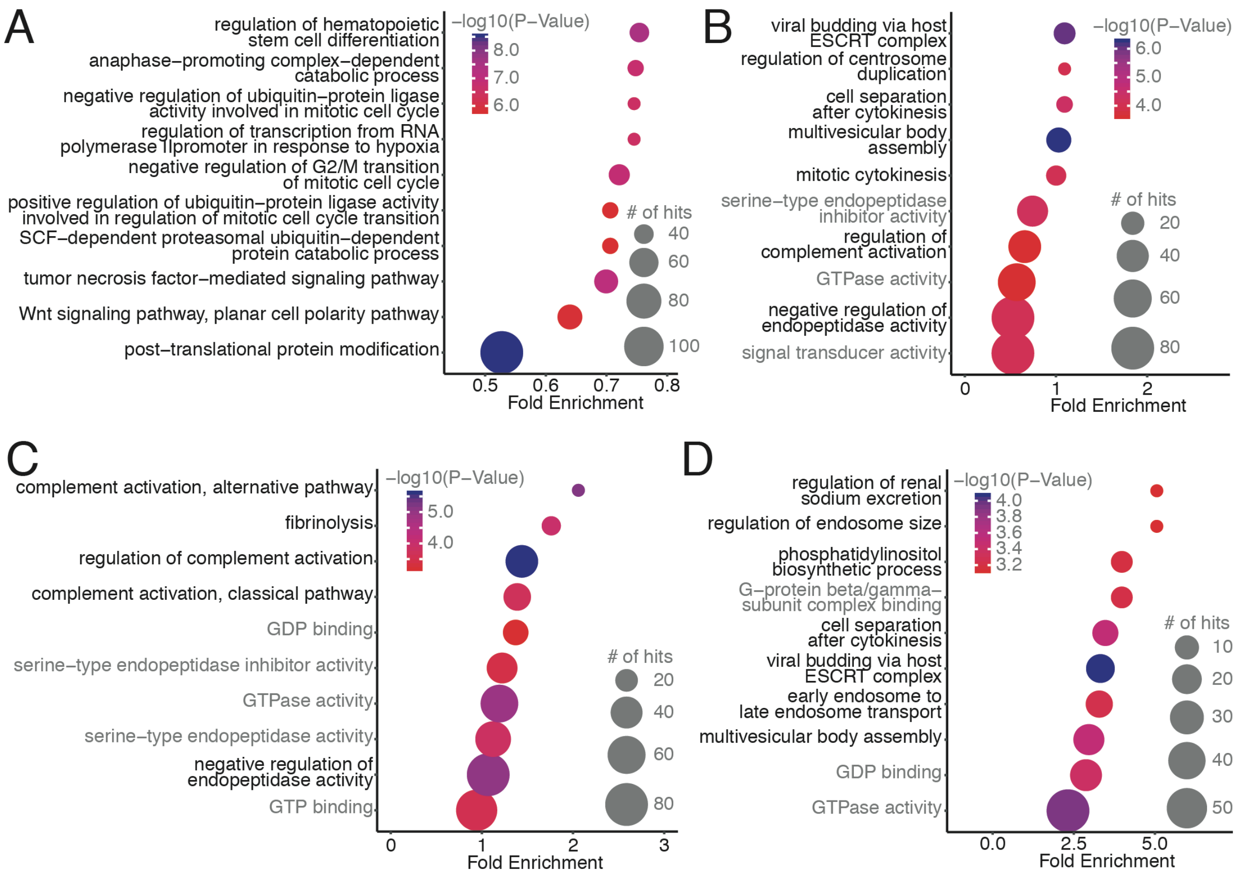


**Fig.S3: Significant GO term analysis of protein fold changes compared to initial donor sample.** Top 10 significant GO terms for each timepoint after transplantation compared to the initial donor samples. P value depicted as color code, number of annotated proteins corresponding to bubble size, black GO terms indicating biological processes, gray GO terms indicating molecular function **(A:** Timepoint B vs A**, B:** Timepoint C vs A**, C:** Timepoint D vs A**, D:** Timepoint E vs A**)**.

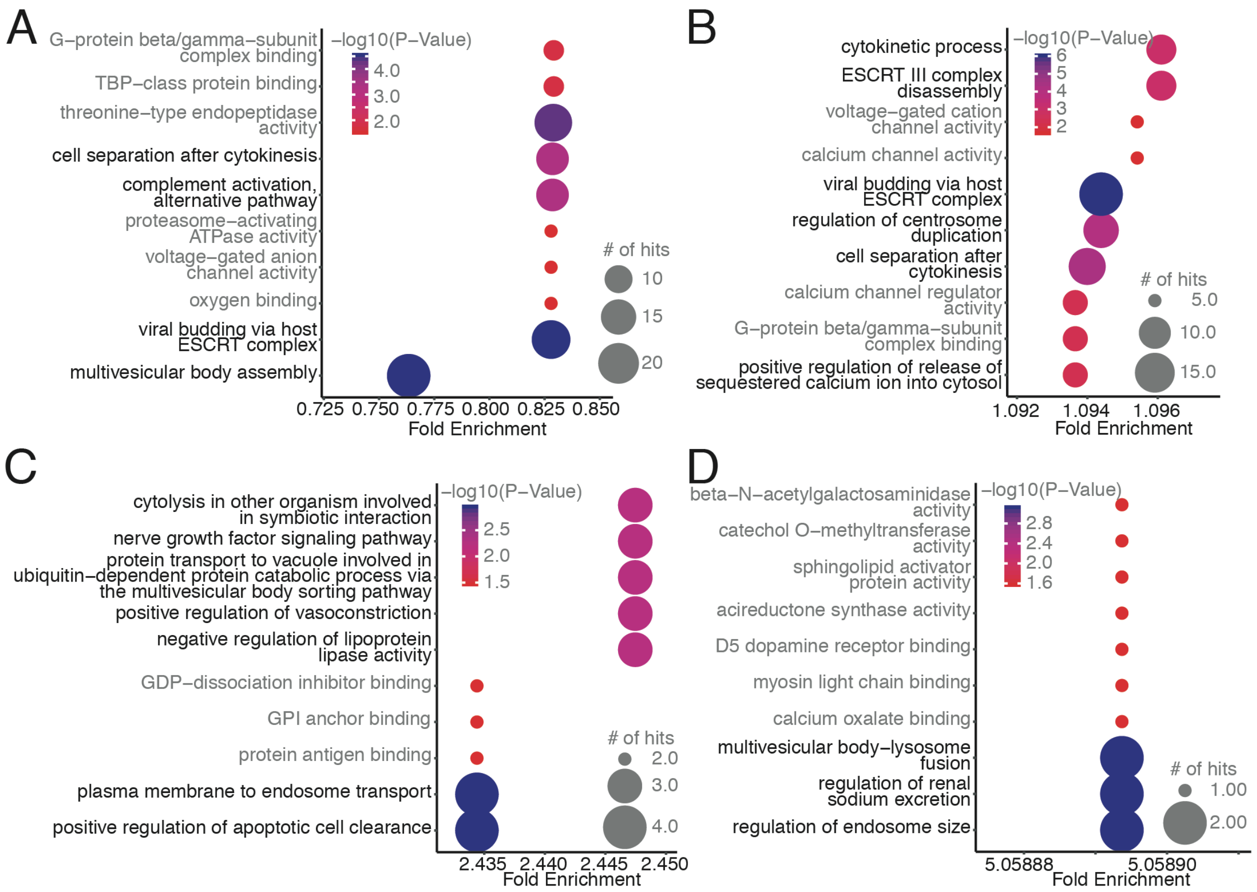


**Fig.S4: Enriched GO term analysis of protein fold changes compared to initial donor sample.** Top 10 GO terms enriched for each timepoint after transplantation compared to the initial donor samples. P value depicted as color code, number of annotated proteins corresponding to bubble size, black GO terms indicating biological processes, gray GO terms indicating molecular function **(A:** Timepoint B vs A**, B:** Timepoint C vs A**, C:** Timepoint D vs A**, D:** Timepoint E vs A**)**.

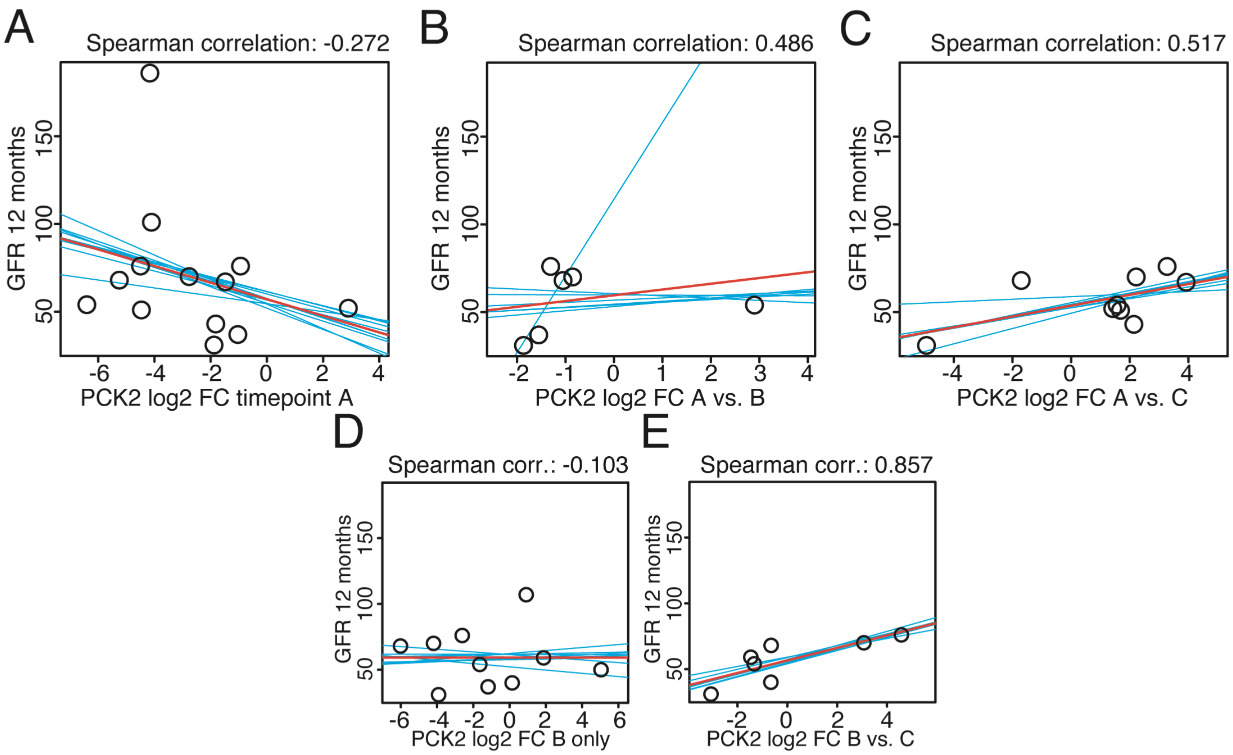


**Fig.S5: PCK2 abundance in suEVs at other timepoints does not correlate with estimated GFR 12 months after transplantation.** Correlation plots of PCK2 intensity foldchange to mean intensity at timepoint A **(A)** and B **(B)** and foldchanges between timepoints A vs. B **(C)**, A vs. C **(D)**, and B vs. C **(E)**, to GFR 12 months after transplantation measured in the validation cohort. Blue: individual linear regression models; Red: merged linear regression model for all samples.

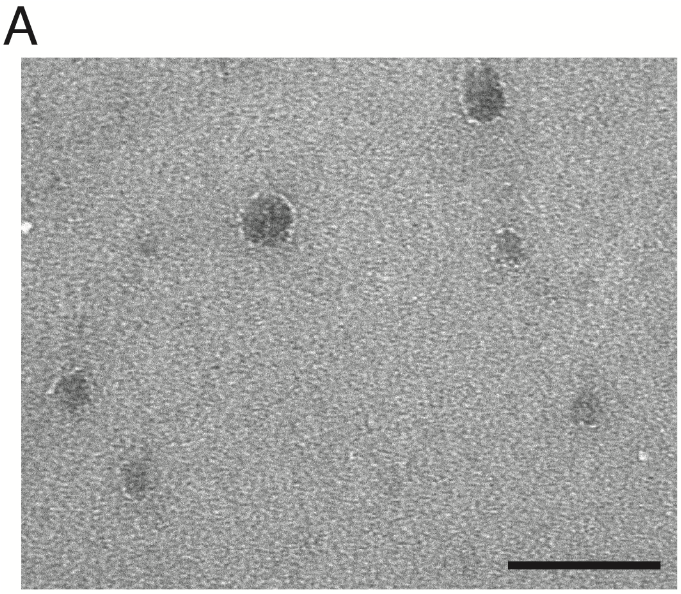


**Fig.S6: PCK2 localizes to suEVs, increases during the initial stages of transplantation and does not correlate to tissue PCK2 levels after reperfusion.** Electron microscopy depicting non permeabilized small urinary extracellular vesicles after immunogold staining for PCK2, scalebar 100nm (A).
